# Anatomy of an Extensively Drug Resistant *Klebsiella pneumoniae* Outbreak in Tuscany, Italy

**DOI:** 10.1101/2021.06.02.446696

**Authors:** Melissa J. Martin, Brendan W. Corey, Filomena Sannio, Lindsey R. Hall, Ulrike MacDonald, Brendan T. Jones, Emma G. Mills, Jason Stam, Rosslyn Maybank, Yoon Kwak, Katharina Schaufler, Karsten Becker, Nils-Olaf Hübner, Stefania Cresti, Giacinta Tordini, Marcello Valassina, Maria Grazia Cusi, Jason W. Bennett, Thomas A. Russo, Patrick T. McGann, Francois Lebreton, Jean-Denis Docquier

## Abstract

A protracted outbreak of New Delhi metallo-beta-lactamase (NDM)-producing carbapenem-resistant *Klebsiella pneumoniae*, started in Tuscany, Italy, in November 2018 and continued in 2020 and through 2021. To understand the regional emergence and transmission dynamics over time, we collected and sequenced the genomes of 117 extensively drug-resistant, NDM-producing *K. pneumoniae* isolates cultured over a 20-month period from 76 patients at several health care facilities in South-East Tuscany. All isolates belonged to high-risk clone ST-147 and were typically non-susceptible to all first line antibiotics. Albeit sporadic, resistances to colistin, tigecycline and fosfomycin were also observed as a result of repeated, independent mutations. Genomic analysis revealed that ST-147 isolates circulating in Tuscany were monophyletic, highly genetically related (including a network of 42 patients from the same hospital and sharing nearly identical isolates) and shared a recent ancestor with clinical isolates from the Middle East. While the *bla*_NDM–1_ gene was carried by an IncFIB-type plasmid, our investigations revealed that the ST-147 lineage from Italy also acquired a hybrid IncH-type plasmid carrying the 16S methyltransferase *armA* gene as well as key virulence biomarkers often found in hypervirulent isolates. This plasmid shared extensive homologies with mosaic plasmids circulating globally including from ST-11 and ST-307 convergent lineages. Phenotypically, the carriage of this hybrid plasmid resulted in increased siderophore production but did not confer virulence to the level of an archetypical, hypervirulent *K. pneumoniae* in a subcutaneous model of infection with immunocompetent CD1 mice. Our findings highlight the importance of performing genomic surveillance to identify emerging threats.

**Significance Statement:** Carbapenem-resistant *Klebsiella pneumoniae* belong to the “critical priority” tier of bacterial pathogens as identified by the World Health Organization. Emerging “high-risk” lineages are responsible for difficult-to-treat, hospital-acquired infections and outbreaks around the globe. By integrating genomic and epidemiological data for isolates collected over 20 months, this study revealed both the high, regional prevalence and the rapid spread, within a single hospital, of *K. pneumoniae* ST-147 in Italy. Besides resistance to nearly all antibiotics, we showed that this lineage carried a hybrid plasmid harboring a set of biomarker genes previously linked to hypervirulence. Convergence of multidrug resistance and hypervirulence is a major concern and these findings highlight the need for robust, global surveillance to monitor the emergence of high-risk *K. pneumoniae*.

## Introduction

*Klebsiella pneumoniae* is a leading cause of healthcare-associated infections including pneumonia, urinary tract, and bloodstream infections (1). These classical *K. pneumoniae* (cKp) frequently cause opportunistic infections in immunocompromised patients, elderly, neonates, and patients with inserted medical devices (2). Of further concern, cKp strains can readily acquire antimicrobial resistances including extended-spectrum betalactamases and carbapenemase-encoding genes (3). Prompting a major public health challenge, the global emergence and dissemination of multidrug resistant *K. pneumoniae* (MDR-cKp) is attributed to a few successful clonal lineages, including newly identified “high-risk” sequence type (ST)-147 and ST-307 (4–6). Similar to ST-307, the MDR-cKp ST-147 lineage emerged in Europe during the mid-1990s, acquired plasmids encoding various carbapenemases genes (*i.e*., VIM- and NDM-type metallo-β-lactamases as well as OXA-48 serine-carbapenemase variants) in the mid-2000’s, and has now spread to all continents (4).

In recent years, a distinct hypervirulent pathotype (hvKp) has been identified, and is recognized clinically by invasive and disseminated infections, in otherwise healthy individuals, that include meningitis, liver abscesses, and endophthalmitis (2). However, the defining features of hypervirulence remain ambiguous. Phenotypically, hvKp isolates have been characterized primarily by their hypermucoviscosity, antimicrobial susceptibility, and greater production of siderophores. Genetically, several virulence genes, carried on large virulence plasmids and integrative conjugative elements (ICE), encoding for the biosynthesis of siderophores (aerobactin [*iuc*], salmochelin [*iro*] and yersiniabactin [*ybt*]), the modulation of mucoviscosity and capsule synthesis (*rmpADC/rmpA2),* metabolite transporter *peg-344,* and the production of genotoxic polyketide colibactin (*clb*) have been linked to the hvKp pathotype (5, 7, 8)

Compounding the problem, convergent *K. pneumoniae* lineages with both virulence and resistance genes have been observed, albeit infrequently (3, 9, 10). Genotypic convergence most frequently occurs when MDR-cKp lineages acquire mobile genetic elements that carry the aforementioned virulence biomarker genes (11). Alarmingly, large hybrid plasmids that harbor both antimicrobial resistance and virulence genes have recently been reported in MDR isolates from multiple countries (6, 12–16). This includes, sporadic instances of convergent *bla*_NDM_-carrying-ST-147 isolates detected in the United Kingdom (U.K.) in 2018 and 2019 (13, 14). Yet, in most cases, the clinical impact and the resulting virulence potential of convergent lineages is not well understood.

In this report, we investigated the emergence, genotypic convergence, phenotypic virulence, as well as the regional and nosocomial spread of a NDM-producing ST-147 clone causing an outbreak in Tuscany, Italy (17). This outbreak caused by an extensively drug-resistant (XDR) *K. pneumoniae* was first identified in November 2018 in the Tuscany region where a significant increase in reported cases from seven hospitals led to expanded surveillance practices (18, 19). Recent surveillance data (Communicable Disease Threats Report 24-30 January 2021 by the European Centre for Disease Prevention and Control) suggest that, at the time of writing, this outbreak is ongoing with the risk of further transmission within and beyond Italy.

## Results

### Emergence of the NDM-1 producing ST-147 outbreak clone in Italy

Between February 2019 and October 2020, a complete collection of NDM-producing *K. pneumoniae* ST-147 from the University Teaching Hospital of Siena and several long-term health care facilities in the South-East (SE) Tuscany region resulted in 117 isolates cultured from 76 patients (44% female; age ranging 27-96; median, 75) (**Table S1**). Similar to ST-147 strains previously reported from the North-West (NW) region of Tuscany (19), these isolates were predominantly non-susceptible to all tested penicillins, cephalosporins (including in combination with a β-lactamase inhibitor), carbapenems, and aminoglycosides (**Table S2**). These 117 isolates were mostly cultured from rectal swabs (74%; though the fraction of asymptomatic carriage is unknown), urine (14%), respiratory tract (6%), or blood (5%) clinical samples. In 4/6 patients (ID# 20, 25, 26, and 40) with bloodstream infections, positive rectal swab preceded blood cultures suggesting gut-colonization resulted in invasive disease (**Table S1**). Whole-genome sequencing (WGS) and phylogenetic analysis revealed that all ST-147 isolates from Italy (including 9 publicly available genomes from bloodstream isolates collected in the NW Tuscany (19)) were monophyletic (node 1) and showed high genetic relatedness (average pairwise distance from nearest neighbor was 5.5 SNPs) (**Fig. 1**).

**Figure 1.**
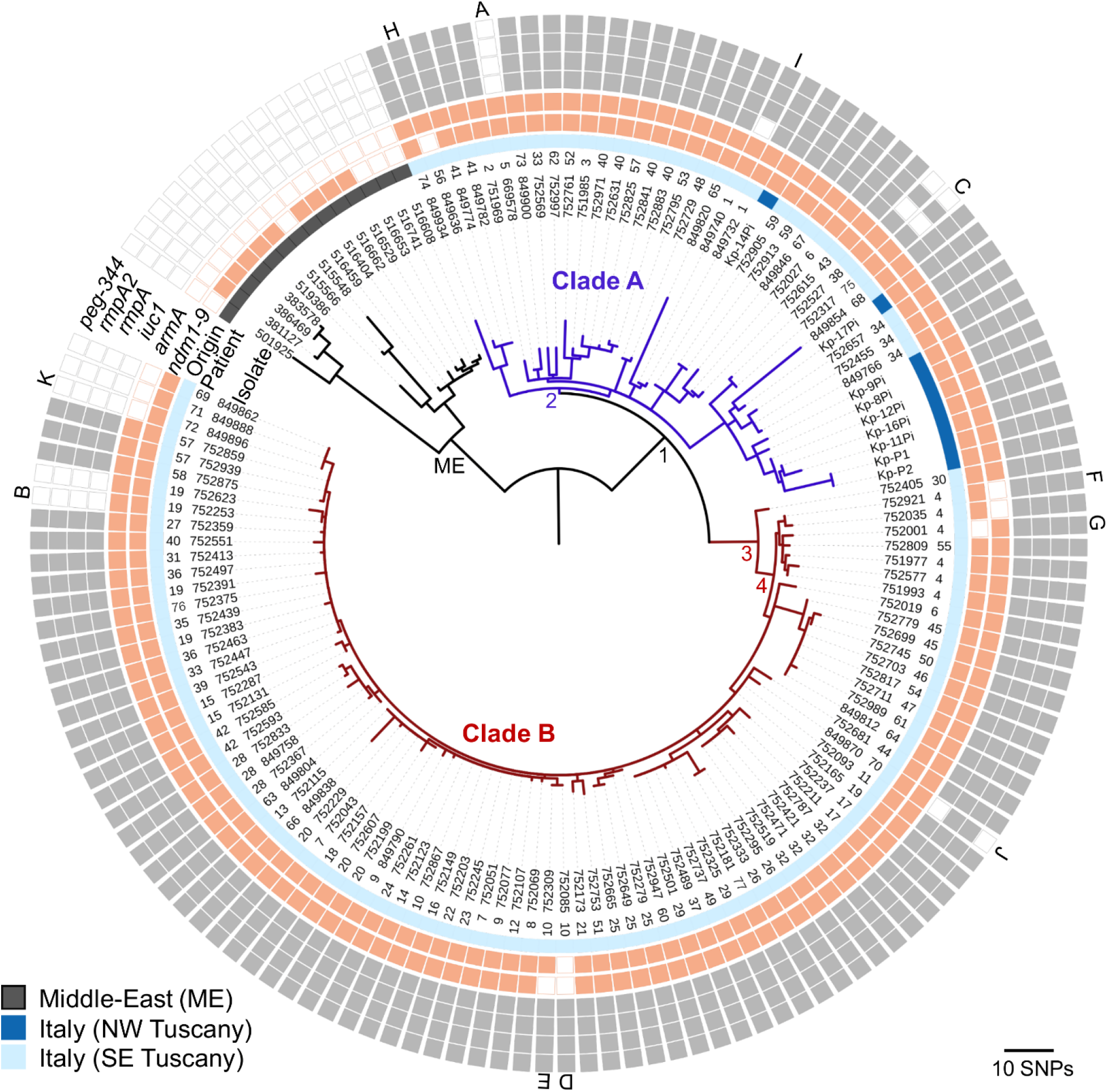
Maximum likelihood core SNP-based phylogeny of *K. pneumoniae* ST-147 from Italy. Closely related isolates from the Middle East were included as an outgroup. Nodes of interest (all with bootstrap values > 80) are labeled (ME, 1 to 4). When available, numerical patient identifiers are provided (1 to 77). Concentric rings indicate the presence (closed symbol) of a selection of plasmid-bound resistance and virulence genes. Apparent variations in plasmid content are labeled (A to K) on the outermost ring.

When compared to international ST-147 genomes (from both public databases and the MRSN collection) the outbreak clone from Italy most closely resembled (shortest distance was 45 SNPs) isolates collected from the Middle East (ME) between 2005 and 2015 (**Fig. 1 and S1**). All isolates were predicted to be K-antigen capsular biosynthesis loci K64 and O-antigen type O2 variant 1 (O2v1) (**Table S1**). Further, the isolates from Italy carried a *bla*_NDM–1_ metallo-β-lactamase gene on an IncFIB-type plasmid (54,064 bp; hereby named pSI0739-NDM) (**Fig S2a)** and a comparable plasmid (99% identity and 87% coverage) was found in 9 of 14 isolates from the ME (**Fig 1**).

### Convergence of virulence and resistance genes

Genomic comparisons further identified that, since divergence from the ancestor shared with the ME lineage, the ST-147 clone from Italy acquired a large hybrid IncH-type plasmid (335,489 bp), subsequently referred to as pSI0739-ARMA-Vir (**Fig S2b**). When compared to public databases, pSI0739-ARMA-Vir was highly related to hybrid plasmids previously characterized from various countries (U.K., Germany, and Czech Republic), and distinct convergent *K. pneumoniae* lineages (ST-307, ST-15, ST-147, and ST-383) (**Table S3**) (6, 14, 20). In addition to 8 other antimicrobial resistance genes (**Fig. S2b**), pSI0739-ARMA-Vir carries the 16S rRNA methyltransferase, *armA*, as well as virulence genes encoding a tellurium resistance operon (*terZABCDEF*), aerobactin biosynthesis (*iuc1*), putative hypermucoidy and capsule synthesis loci (*rmpADC*/*rmpA2*), and the metabolite transporter, *peg-344* (**Fig. 1 and S2b**). 92% of isolates also carry genes encoding the yersiniabactin siderophore system (*ybt9* lineage harbored by a chromosomal ICEKp3).

Within the outbreak isolates from Tuscany, the presence or absence of plasmid-bound resistance and virulence genes was used to identify excision events or plasmid loss (labelled A-K) (**Fig. 1 and S2b)**. In four independent instances (A, B, I, and J) a ~ 60 to 90 Kb segment containing the virulence genes was excised from pSI0739-ARMA-Vir (**Fig. 1 and S2b**). Insertion sequences (IS) were often identified at the boundaries of the virulence island, suggesting transposition could play a role in the mosaic structure of these plasmids, as previously proposed (14). Notably, excision event B occurred within the host where (2 out of 5) serial isolates from patient 19 eventually lost all virulence genes (**Fig. 1 and S2b**). The *armA* gene cassette, flanked by IS*26* elements, was also excised on three separate occasions (D, E and F; the latter as the likely result of in-host evolution within serial isolates from patient 4). By contrast, the complete pSI0739-ARMA-Vir was lost in three of the later isolates (July and August 2020; event K); all highly genetically related but from distinct patients (**Fig. 1 and S2b**). Plasmid loss was also observed for pSI0739-NDM (three distinct events D, G, and H) (**Fig. S2b**). As expected, susceptibility to the aminoglycosides and/or carbapenems was restored in these isolates (**Table S2**).

### ST-147 outbreak is fueled by nosocomial spread

Phylogenetic analysis of the isolates from Tuscany revealed the existence of two clades (differing by 32.9 SNPs on average), A and B (**Fig. 1**). Clade A was relatively heterogenous (average pairwise distance from nearest neighbor was 11.1 SNPs) and comprised 31 isolates from 23 patients at 3 distinct healthcare facilities in SE Tuscany (**Fig. 1 and Table S1)**. Additionally, clade A includes 9 genomes from a hospital in NW Tuscany for which patient metadata is unknown (19). In contrast, clade B was highly homogeneous (average distance of 2.4 SNPs from nearest neighbor) and grouped 86 isolates collected from 57 inpatients from the Siena University Teaching hospital.

Temporally, detection of the first clade A isolate from each patient (*i.e.* removing serial isolates) was relatively stable (1.2 new cases per month on average) across the 20-month sampling period (**Fig. 2a**). By contrast, a large increase of new cases due to clade B isolates (7.1 cases per month) was observed over a 6-month period (July to December 2019). Further, the majority (90%) of clade B isolates were collected from only 5 wards with few patients (n = 10) yielding positive, serial cultures from multiple wards **(Fig. 2b)**. To trace possible nosocomial events, patient pairs that shared an isolate separated by 4 SNPs or less were identified and used to construct a network of possible transmission. This approach ultimately linked 74% (n = 42) of all patients represented in clade B and revealed that 8 inpatients (ID# 7, 9, 10, 14, 18, 20, 22, and 23), all in the cardiology clinic in July and August of 2019, shared genetically identical isolates (**Fig. 2bc**). While the last case detected in the cardiology clinic was in September 2019, this clone had seemingly spread throughout the hospital. Indeed, between September 2019 and October 2020, the internal medicine and emergency wards accounted for 75% of primary cases due to a clade B isolate (**Fig. 2bc**). Of note, the apparent reduction in isolates collected from patients between February and July 2020 was likely the result of hospital policies to reduce the number of non-urgent programmed hospitalizations in relation to the COVID-19 pandemic. New cases (*e.g*. patients 69-72) quickly reappeared in the fall of 2020 indicating nosocomial transmission of clade B was not yet contained (**Fig. 2ab**).

**Fig. 2.**
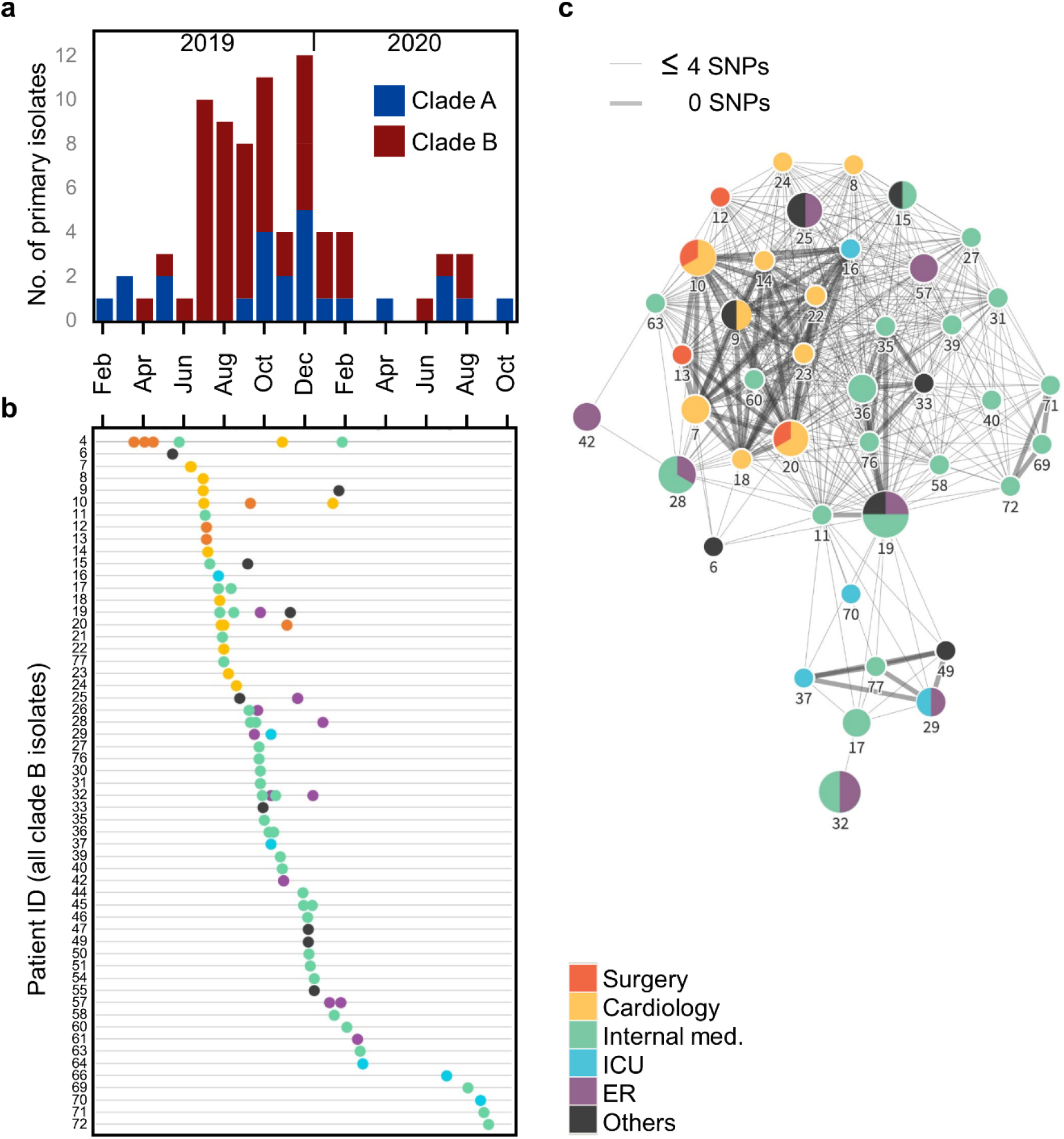
Rapid, nosocomial spread of Clade B ST-147 clone. **(A)** Epidemic curve of all patient’s first isolate (i.e. no serial isolates) collected in SE Tuscany and colored by phylogenetic clade (A or B). **(B)** Patient charts showing the date (x-axis) and location (colored legend) of all first and serial clade B isolates (dots). **(C)** Network of patients with near-identical clade B isolates (≤4 SNPs). Nodes are labeled with their respective patient ID and indicate the location (colored pie chart) and number (size) of serial isolates.

### Positive selection on resistance and virulence determinants

In addition to some isolates independently excising regions within or the whole pSI0739-ARMA-Vir and pSI0739-NDM plasmids, mutational convergence was also observed within specific genes and functional clusters. The most striking example was gene *glpT* that encodes the main transporter responsible for the uptake of the fosfomycin antibiotic. Not accounting for synonymous (SYN) mutations, 10 variants emerged independently in *glpT* (4 predicted loss-of-function [LOF] and 6 non-synonymous [NSY] mutations) and were found in 24 outbreak isolates, mostly from clade A (**Fig. 3**). This is in agreement with a previous report of *glpT* mutations found in clinical, epidemic ST-147 isolates that was likely responsible for reduced fosfomycin susceptibility (21). Four distinct NSY mutations in the *cusSRA* cluster, previously associated with copper and silver, and tigecycline resistance (22), were found in 6 isolates from both clades (**Fig. 3**). In addition, the *ramR-ramA* gene cluster, consisting of the *romA-ramA* operon and the divergently transcribed *ramR* (resistance antibiotic multiple protein A), was mutated at 7 distinct occurrences (including 2 predicted LOF in *ramR* and one in *romA*) in 20 strains (**Fig. 3**). Two serial isolates from patient 4 with the same frameshift mutation in *ramR* demonstrated tigecycline resistance (MIC ≥ 8 mg/L), consistent with previous reports in *K. pneumoniae* (23) (**Fig. S4b and Table S2**). Notably, isolate 752623 also evolved tigecycline resistance, but the genetic determinants remain unclear (**Table S2**). The other predicted LOF mutations in both *ramR* and *romA* were found in 5 genetically related isolates from NW Tuscany (Kp-P1, Kp-P2, Kp-11Pi, Kp-12Pi and Kp-16Pi) for which decreased susceptibility to tigecycline was noted (19). Interestingly, Falcone *et al*. also reported that: i) isolates Kp-P1 and Kp-P2 carried *bla*_NDM–9,_ which differs from *bla*_NDM–1_ by a single amino acid substitution and confers higher levels of resistance to the carbapenems, and ii) isolates Kp-P1, Kp-P2, Kp-12Pi, and Kp-16Pi evolved resistance to colistin in part due to loss of function mutations in *mgrB* (19). Here, isolate 849820 carried a truncated *mgrB* (**Table S1**) but remained susceptible to colistin (MIC < 0.25 mg/L). Positive selection was also observed for genes involved in cell wall synthesis (*pbp2*, *pbp3*, *rlpA*) and glycosylation (*pglA*), adhesion to mucosal surfaces and virulence (*fimH*), nitrate respiration (*narLXG*), and iron binding and transport (*sufBDS*) (**Fig. 3**). The latter was recently reported as a potential factor underlying the persistence of an outbreak involving high-risk ST-258 *K. pneumoniae* (24).

**Fig. 3.**
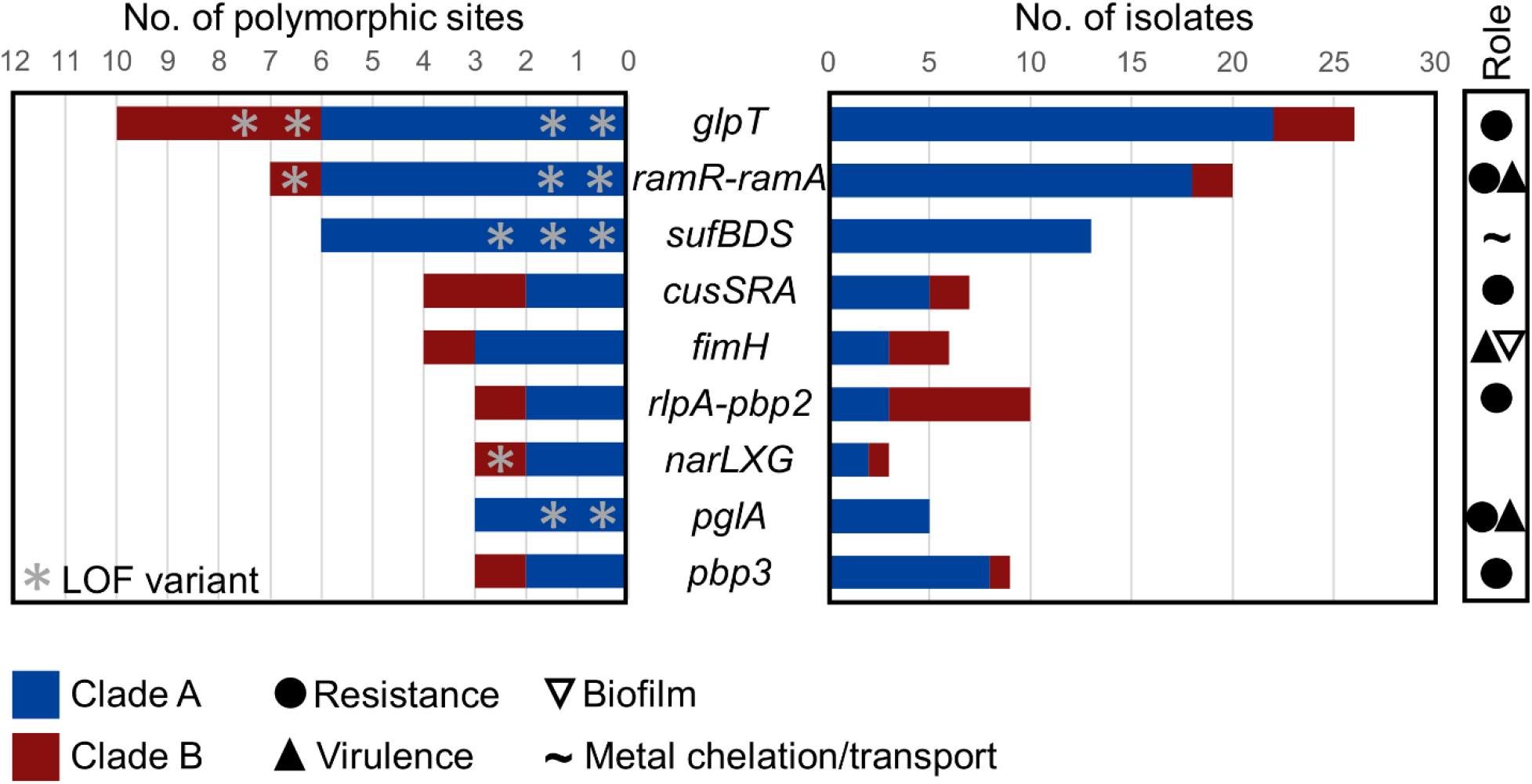
Convergent evolution on resistance and virulence genes. List of genes (or functional clusters of syntenic genes) that were repeatedly and independently (3 or more) mutated among the outbreak isolates. Bars show the number of different mutations within each gene or functional cluster (left diagram) and the numbers of isolates with a mutated allele (right diagram). Data is plotted for isolates from both phylogenetic clades. When available, and based on the literature, the role of genes and functional clusters in relevant phenotypes is indicated.

### Fixation of variants through patient transmission and within host evolution

To further identify traits that possibly contributed to the success and spread of the ST-147 clone in hospitals in Tuscany, we identified variants that became fixed in the population at nodes of interest (excluding SYN and intergenic variants) (**Fig. S3a**). Using parsimony, we identified 8 variants found in all isolates from Italy that were missing in all ME genomes (node 1). Of particular interest were: (i) a predicted LOF mutation in *cycA* which encodes a Serine/Alanine transporter that, when functional, allows the antibiotic D-cycloserine to cross the cell wall and reach its cytoplasmic target (25), (ii) a NSY mutation in *pabA* involved in aromatic amino acid biosynthesis and previously characterized as a general infection requirement for *K. pneumoniae* (26), and (iii) a NSY mutation in *hdfR* encoding for a transcriptional regulator involved in the complex regulation of capsule production and hypermucoviscosity (27).

Two NSY variants (in genes *fliY* and *ybiO*) arose in the last common ancestor at the root of clade A (node 2) while a total of 9 variants arose at the root (nodes 3 and 4) of clade B (**Fig. 1 and S3a**). Among them, variants in two genes could participate in the apparent success of clade B in spreading and persisting in the hospital: *metQ* a D-methionine-binding lipoprotein, shown to be important for colonization (28), and *pal*, encoding a peptidoglycan-associated lipoprotein shown to provide protection against neutrophil phagocytosis and killing as well resistance to gastrointestinal bile salts (29), a key ability for transmission via the fecal-oral route. Similarly, longitudinal sampling of six serial isolates (March 2019 to January 2020) from patient 4 revealed a stepwise, fixation of variants (**Fig. S3b**). NSY mutations were present in genes involved in cell wall biosynthesis/recycling (*rlpA*, *emtA*) and surface polysaccharide synthesis (*rffG*). The latter is part of a gene cluster that encodes the Enterobacterial Common Antigen (ECA), a carbohydrate polymer thought to play a significant role in *K. pneumoniae* physiology and host interactions (26). Amino acid substitutions in the adhesin *fimH* and in the multifunctional regulator *mgrA*, were also observed in these serial isolates.

### Pathogenicity of the convergent ST-147 clone from Italy

While sequencing allowed for the detection of genotypic convergence, the virulence of this epidemic ST-147 clone remained to be established. Two distinct outbreak isolates carrying pSI0739-ARMA-Vir (752019 and 752165) produced siderophore at levels well above the 30 μg/ml threshold predictive of the hvKp pathotype (~230 μg/ml**; Fig. 4a and S4a**) (30). The excision of virulence genes (including the *iuc* siderophore biosynthesis operon) in outbreak isolate 752253 (event B, **Fig. S2b**) resulted in siderophore levels comparable to the cKP1 control (**Fig. S4a**). However, despite carrying the virulence genes *rmpADC* and *rmpA2*, isolates 752019 and 752165 showed mucoviscosity levels comparable to the cKP1 control (**Fig. 4b and S4b**). Moreover, a subcutaneous (SQ) model of infection with immunocompetent CD1 mice determined that 752019 and 752165 were non-lethal at a challenge inoculum of 10^3^ CFU, unlike canonical hypervirulent isolate hvKP2 (**Fig. 4c**). Further, no lethality was observed with a 10^5^ CFU inoculum of the ST-147 isolates and only SQ challenges with 10^7^ and 10^8^ CFU revealed a slight increase in lethality (non-significant for 752019 [**Fig. 4c**]; *p*=0.029 for 752165 at 10^8^ CFU [**Fig. S5**]) when compared to the cKP1 control, which was consistently found as non-lethal at a challenge inoculum up to 2.5 × 10^8^ CFU (31). Likewise, SQ challenge with strain PBIO1953 (6), a convergent ST-307 *K. pneumoniae* outbreak clone carrying a hybrid plasmid with extensive similarity pSI0739-ARMA-Vir (**Table S3**), resulted in mortality rate of 20% at the highest inoculum (not reaching statistical significance) compared to the cKP1 control (**Fig. S5**).

**Fig. 4.**
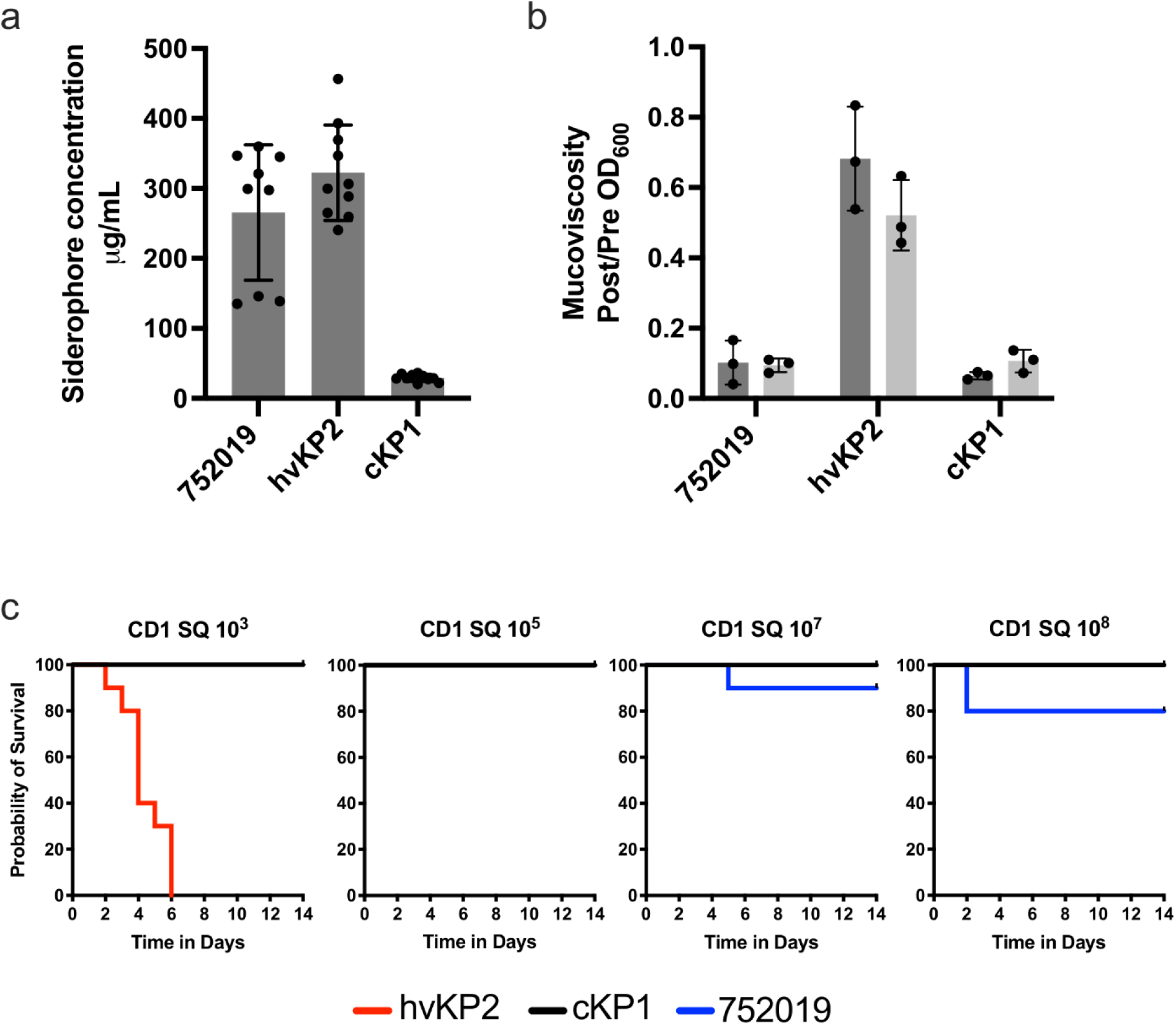
Pathogenicity of the convergent ST-147 clone from Italy. **(A)** Quantitative siderophore production of outbreak isolate 752019 carrying the canonical pSI0739-ARMA-Vir. Reference strains hvKP2 (hypervirulent pathotype) and cKP1 (classical pathotype) are shown throughout all panels for comparison. **(B)** Mean mucoviscosity of outbreak isolate 752019 compared to reference *K. pneumoniae* isolates. Independent assays were performed for both media (LB and M9). **(C)** Kaplan-Meier survival curves of outbred CD1 mice after subcutaneous (SQ) challenge with 10^3^, 10^5^, 10^7^, or 10^8^ CFU of outbreak isolate 752019 or reference isolates hvKP2 and cKP1. Total *n*=10 (*n*=5 in each of 2 independent experiments) for each titer for each strain.

## Discussion

We report a comprehensive analysis of an NDM-producing ST-147 *K. pneumoniae* outbreak clone circulating in Tuscany, Italy. Although the number of colonized/infected patients was significantly lower than in the North-West region (18), our study demonstrates that this epidemic, XDR clone has now spread throughout multiple healthcare facilities in the South-East region of Tuscany. Phylogenetic analysis suggests that this clone possibly emerged from a NDM-1-producing, ST-147, MDR-cKp lineage, previously shown to be endemic in the Middle East (32), by acquiring a hybrid resistance/virulence plasmid. Furthermore, we also confirm the early concern about cross-border spread (17), as highly genetically related isolates (**Fig. S1**), with near identical plasmids (**Table S3**), have been reported in multiple U.K. hospitals in 2018 and 2019 (14).

Signs of strong selective pressure was observed in genes associated with drug resistance. With the prototypical ST-147 outbreak isolate resisting to all first line agents (*i.e*. beta-lactams, including carbapenems; fluoroquinolones; aminoglycosides) (**Table S2**), effective treatment relies on second line antibiotics (*e.g.* colistin, tigecycline and fosfomycin) as well as combination therapies (33). In this context, the repeated, independent emergence of variants associated with resistance to either fosfomycin (*glpT*), tigecycline (*ramR*), and colistin (*mgrB*) is particularly worrisome and raises the concern of the emergence of a pan-resistant ST-147 epidemic lineage.

While WGS allows for the detection of convergence events in *K. pneumoniae*, the definitive impact on pathogenicity often remains unclear (3, 31). Compared to this ST-147 clone from Italy, the acquisition of the same combination of virulence genes was reported in distinct lineages of *K. pneumoniae* ST-11 from China and Egypt (15, 16), and ST-307 from Germany (6). Comparable virulence was observed in a *Galleria* insect model at 10^6^ CFU of ST-11 strains with and without the virulence plasmid (15). Furthermore, using clinical cohort stratifying, no increased mortality was observed in patients infected with ST-11 isolates, carrying the virulence plasmid, (15). The recent ST-307 epidemic clone from Germany carried a near identical hybrid plasmid to pSI0739-ARMA-Vir (**Table S3**) and similarly showed increased siderophore production directly tied to the presence of the virulence genes (6). Here, a SQ challenge of immunocompetent CD1 mice unequivocally demonstrates that isolates from both lineages (ST-147 from Italy and ST-307 from Germany) did not exhibit a maximal hypervirulent phenotype (where lethality is observed with < 10^3^ CFU (31)) and were only slightly more lethal than a cKP, at the highest inoculum tested. Nonetheless, a small increase in pathogenic potential may be clinically significant, particularly in the healthcare setting in which patients are variably immunocompromised.

Based on the current knowledge of *K. pneumoniae* pathogenicity, the reason(s) for the minimal virulence of the convergent ST-147 and ST-307 outbreak clones remains unclear. While increased siderophore production was historically associated with hvKp isolates (34, 35), we have recently shown that this measurement only partially correlates with virulence differences *in vivo* (31). The same observation was made for hypermucoviscosity (31), which, despite the presence of the required *rmpADC*/*rmpA2* genes, was not observed in this ST-147 clone. Other possible reasons are the absence of other known virulence factors (*e.g*. salmochelin and colibactin), genes whose role in virulence has yet to be recognized, virulence gene polymorphism, and differences in capsule type. In this latter possibility, ST-147 isolates are K-antigen type 64, unlike the highly serum resistant K1 and K2 capsules often associated with hvKp isolates (3). Finally, tight regulation of virulence determinants could mitigate the potential biological cost of expressing the many genes on these large hybrid plasmids, a barrier previously proposed as a reason for the relative rarity of convergent strains (3).

The role played by this IncH hybrid plasmid, and the selective forces underlying its emergence in distinct MDR *K. pneumoniae* lineages, remains a central question. While the presence of antibiotic resistance genes undoubtedly confers a selective advantage (36), a hypothesis would be that acquisition of other genes carried by this IncH hybrid plasmid confer increased colonization or persistence abilities, traits not examined in our systemic infection model. This could bear clinical significance as high loads of *K. pneumoniae* in the GI tract have been associated with elevated risk of bacteremia (37). In our collection, most ST-147 isolates were recovered from rectal swabs. Previous work did demonstrate that MDR-cKp colonized poorly in comparison to hvKP1, suggesting an advantage might exist for carrying the virulence plasmid in the gut (38). In fact, increased siderophore production, the one feature observed in ST-147 and a direct result of the presence of the hybrid plasmid, has already been proposed to facilitate gut colonization and spread of various pathogenic *Enterobacterales* (39, 40). Finally, a tellurium resistance operon (*terZABCDEF*), harbored by the IncH hybrid plasmid in both ST-147 and ST-307 outbreak clones, has recently been proposed as a microbiome-dependent fitness factor involved in gut colonization in a murine model (41).

In summary, we identified a convergent clone of ST-147 that shares a recent common ancestor with isolates collected in the Middle East and has been sporadically detected in multiple hospitals in the U.K.. While this outbreak clone was XDR, the maximal hypervirulent phenotype was not observed in immunocompetent hosts *in vivo* despite the presence of a hybrid resistance/virulence plasmid. Further studies are needed to fully characterize the role of these plasmids and the drivers of their emergence in distinct lineages of MDR-Kp around the globe. Close monitoring of global *K. pneumoniae* populations, as well as ongoing, local efforts to contain the dissemination of such high-risk clones are critical to impede the emergence of this troublesome pathogen.

## Materials and Methods

### Bacterial Isolates and Antibiotic Susceptibility Testing

The complete set of XDR *K. pneumoniae* isolates (**Table S1**) were collected from the University Teaching Hospital of Siena (Azienda Ospedaliera Universitaria Senese AOUS), Italy, a 700-bed regional health care facility providing microbiological analyses in the frame of a regional surveillance network for several other local healthcare facilities part of the South-East district and long-term healthcare structures. Bacterial identification and antimicrobial susceptibility testing were carried out using the MALDI Biotyper (Bruker Daltonics GmbH, Bremen, Germany) and the Becton Dickinson Phoenix M50 (Eysins, Switzerland). Carbapenemase production was confirmed using the mSuperCARBA medium (ChromAgar, Paris, France). β-lactamase genes were identified using a PCR analysis (Cica Geneus ESBL Genotype Detection kit, Cica Geneus AmpC Genotype Detection kit and Cica Geneus Carbapenemase Genotype Detection kit 2, Kanto Chemical Co., Tokyo, Japan), as recommended by the manufacturer. Isolates were sent to the Multidrug Resistant Organism Repository and Surveillance Network (MRSN) for further phenotypic characterization and genome sequencing. Confirmatory antibiotic susceptibility testing (AST) was performed in the MRSN College of American Pathologists (CAP)-accredited clinical lab using the Vitek 2 (card GN AST 71 and GN ID; bioMérieux, NC, USA) (**Table S2**). MICs of colistin were determined in triplicate using the broth microdilution method following Clinical and Laboratory Standards Institute guidelines.

### Whole Genome Sequencing

Genomic DNA was extracted and sequenced via Illumina MiSeq or NextSeq benchtop sequencer (Illumina, Inc., San Diego, CA) as previously described (42). For isolate 752165 (alternate name SI-0739) long read sequencing was carried out using a MinION sequencer (Oxford Nanopore Technologies). Basecalling was performed using Guppy (configuration r9.4.1_450bps_hac), filtered using Filtlong (https://github.com/rrwick/Filtlong) and hybrid assembly was performed using Unicycler (43). Complete plasmids, pSI0739-ARMA-Vir and pSI0739-NDM, from isolate SI-0739 were used as reference for plasmid comparison. All genomes have been deposited in the National Center for Biotechnology Information under BioProject PRJNA725484.

### Bioinformatic analysis

Species identification, MLST typing, virulence locus, capsule (K), and lipopolysaccharide (O) loci were identified using Kleborate v2.0.1 (**Table S1**) (44). The *peg-344* gene was identified using BLASTn search of draft genome assemblies (query sequence pLVPK, accession number NC_005249). AMRFinderPlus v3.9.8 (45) and ARIBA v2.14.4 (46) were used to identify resistance alleles from draft assemblies and processed reads, respectively, followed by deduplication of redundant alleles calls. cgMLST allele assignment and minimum spanning tree generation were performed with SeqSphere (47).

We created a SNP phylogeny for ST-147 including the 117 outbreak isolates, 9 previously published genomes (19), and 14 MRSN genomes from ME and Germany (**Table S1**). SNP calling was performed with Snippy v.4.4.5 (https://github.com/tseemann/snippy) using error corrected [Pilon v1.23 (48)] and annotated [Prokka v1.14.6 (49)] draft assembly of 752019 as the reference (**Table S4**). The core SNP alignment was filtered for recombination using Gubbins v2.4.1 (50), which identified 587 variant sites. A maximum likelihood tree was inferred with RAxML-NG v1.0.1 (51) using GTR+G (50 parsimony, 50 random) and was midpoint rooted in iTOL v. 5.5 (52) for visualization with metadata.

For the patient transmission network, a pairwise SNP matrix for clade B isolates was obtained using Snippy v.4.4.5 and snp-dist v.0.7.0 (https://github.com/treemann/snp-dists). Distances ≤ 4 SNPs between isolates from distinct patients were identified. In cases of serial isolates with varying distances, only the lowest inter-patient SNP distance was retained. The resulting network of patients was visualized and edited using Flourish (https://flourish.studio/).

Plasmid content comparisons were performed using closed plasmids pSI0739-NDM and pSI0739-ARMA-Vir as a reference. The draft genomes of outbreak isolates were mapped against the closed reference plasmids using default parameters of the CGView server (http://cgview.ca/).

#### Mucoviscosity and quantitative siderophore assays

Mucoviscosity of selected isolates was measured (ratio of OD_600_ values post/pre centrifugation at 1,000 × g for 5 minutes) after 24 hours of growth (37°C and 275 rpm) in LB or c-M9-te minimal medium (M9), as described previously (31).

To quantify siderophore production, the isolates of interest were grown overnight at 37°C in iron-chelated M9 minimal media containing casamino acids (c-M9-CA) and culture supernatants were assessed using the chromeazurol S dye assay as described (35). For both *in vitro* essays, three independent biological replicates were performed on different days for each strain and the data was reported as the mean ± the SD.

#### Mouse subcutaneous (SQ) infection model

Animal studies were reviewed and approved by the Veterans Administration Institutional Animal Care Committee and the University at Buffalo-SUNY and were carried out in strict accordance with the recommendations in the guidelines delineated in the “NIH Guide for the Care and Use of Laboratory Animals”(revised 1985) and the “Ethics of Animal Experimentation Statement” (Canadian Council on Animal Care, July, 1980) as monitored by the Institutional Animal Care and Use Committee. All efforts were made to minimize suffering. Veterinary care for the animals was supplied by the staff of Veterans Administration Animal Facility under the direction of a fully licensed veterinarian. CD1 male mice, 4-6 weeks old, were obtained from Charles River Laboratories, quarantined for 5 days before use, and challenged via a SQ injection with the isolates of interest (100 μL of bacterial suspension serially diluted to the required titers in 1 × PBS diluted and injected using a 0.5 mL insulin syringe), as previously described (31). The animals were closely monitored for 14 days after challenge for the development of the study endpoints, survival or severe illness (in extremis state)/death, which was recorded as a dichotomous variable. Signs that were monitored and which resulted in immediate euthanasia using methods consistent with the recommendations of the American Veterinary Medical Association Guidelines included hunched posture, ruffled fur, labored breathing, reluctance to move, photophobia, and dehydration

#### Statistical analyses

Statistical analyses were performed using GraphPad Prism 8.0. A Log-rank (Mantel-Cox) test was performed to analyze *in vivo* infection model data. A Mann-Whitney test was used to compare siderophore concentrations between strains. Bonferroni posttests were used to account for multiple comparisons. A *P* value < 0.05 was considered statistically significant.

## Acknowledgments

Thanks are due to Dr. Lerner (ChromAgar, Paris, France) and to Dr. Kawagushi, Dr. Kobayashi and Dr. Yamazaki (Kanto Chemical) for helpful discussions and for kindly providing some of the reagents used in this study.

This study was funded by the U.S. Army Medical command and the Defense Medical Research and Development Program. The manuscript has been reviewed by the Walter Reed Army Institute of Research. There is no objection to its presentation. The opinion or assertions contained herein are the private views of the Department of the Army or the Department of Defense.

This work was also supported by NIH AI123558-01 and 1R21AI141826-01A1 (T.A. Russo) and the Department of Veteran Affairs VA Merit Review (I01 BX004677-01) (T.A. Russo). The funders have no role in the decision to publish or the preparation of this article.

## Supplemental information

**Table S1.** Characteristics of all isolates used in this study.

**Table S2.** Antibiotic susceptibility profiles for a selection of ST-147 isolates.

**Table S3.** Overview of plasmid relatedness with pSI0739-ARMA-Vir..

**Table S4.** List of predicted variants in ST-147 isolates used in this study.

**Figure S1.**
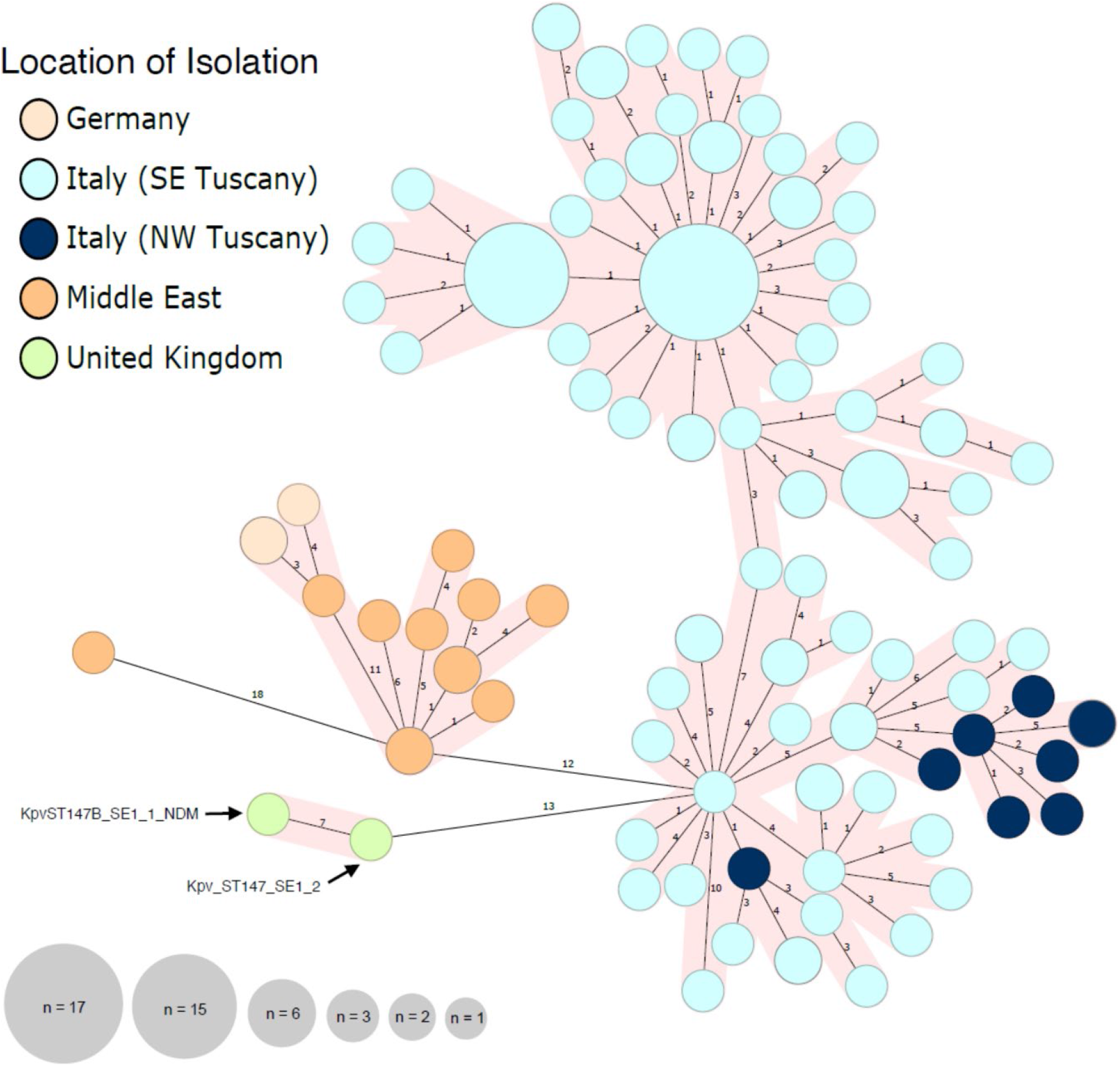
cgMLST minimum spanning tree of 143 ST-147 *K. pneumoniae* genomes used to determine the genetic relatedness of the Italy ST-147 outbreak clone to available ST-147 genomes from the MRSN collection and public databases. cgMLST allelic profiles are represented by circles, and circle sizes are proportional to the number of isolates sharing that profile. The line length connection the isolates represents the number of allelic differences between each profile. Isolates with 11 or less allelic differences are highlighted with pink shading around them.

**Figure S2.**
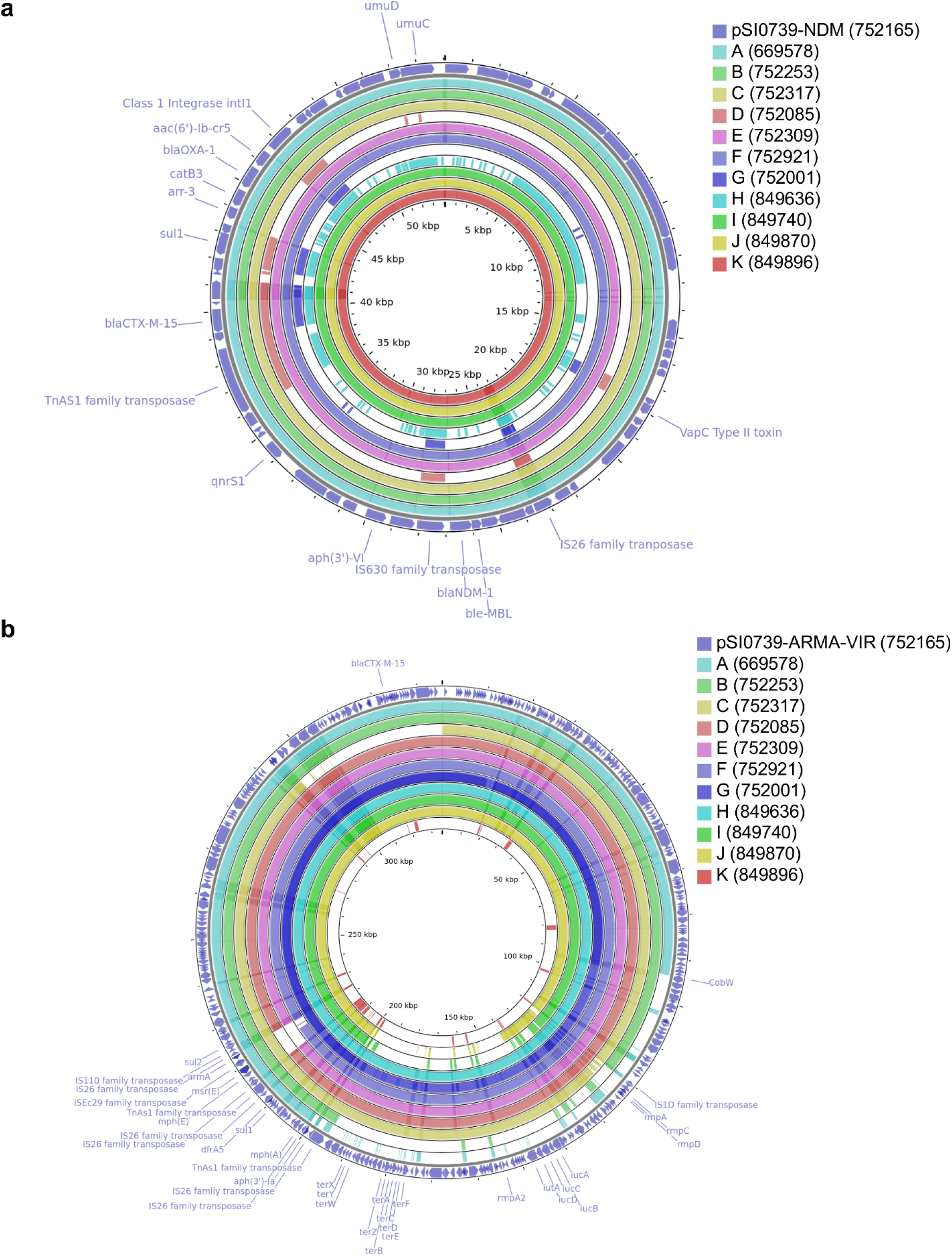
Sequence similarity of hybrid plasmid types identified in this study. **(A)** Sequence assembly contigs from each isolate were blasted against the closed ST-147 NDM carrying plasmid from this study, pSI0739-NDM and **(B)** the hybrid plasmid pSI0739-ARMA-Vir from this study. For each panel, the outer ring represents the reference plasmid. Inner rings represent predicted excision events or plasmid loss identified in a selection (labeled A-K) of outbreak isolates.

**Figure S3.**
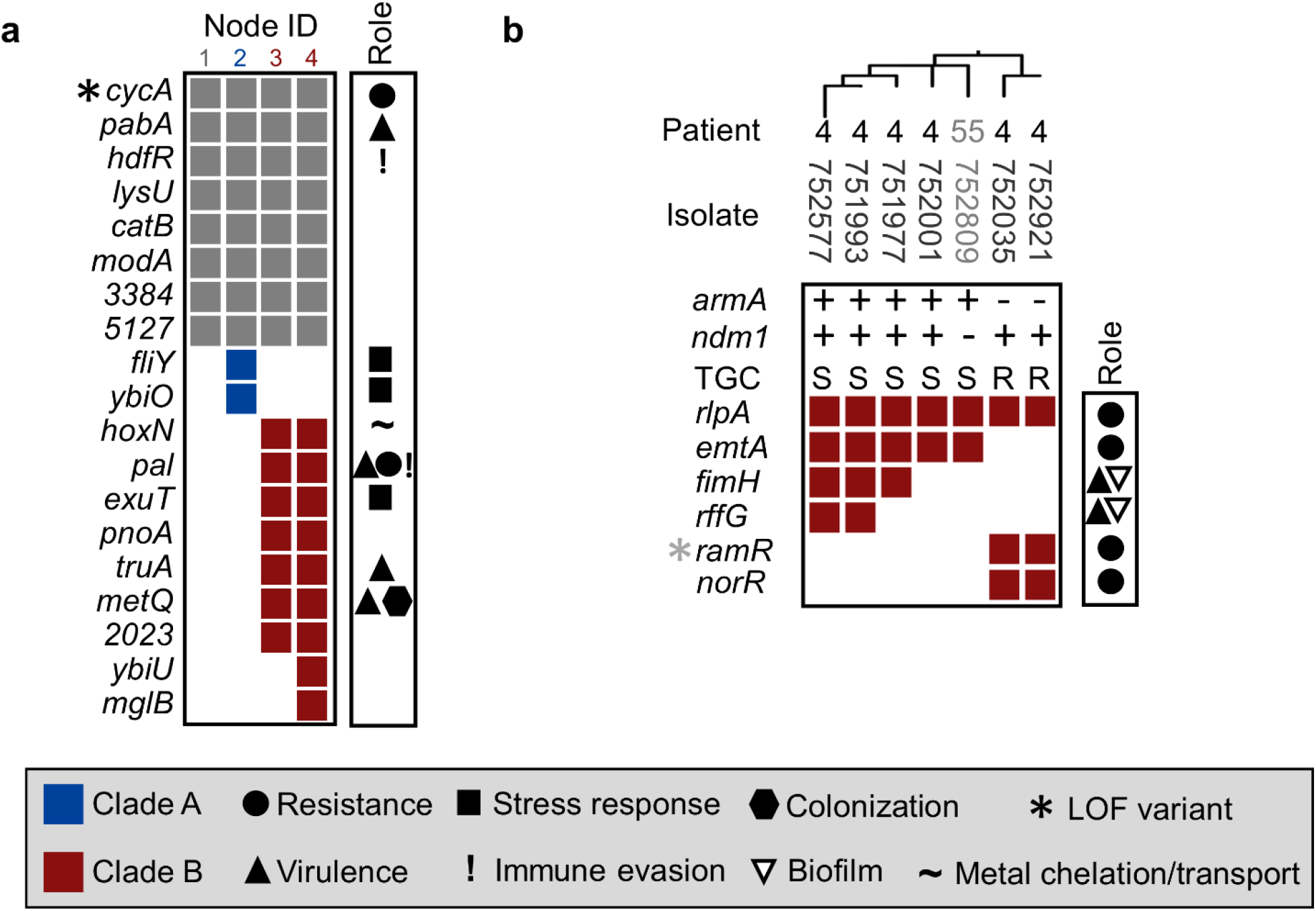
**(A)** Fixation of shared variants within the *K. pneumoniae* ST-147 outbreak strains (nodes 1-4 corresponding to Fig. 1). **(B)** Stepwise, fixation of variants found in 6 serial isolates from patient 4, clade B. For each gene, based on analysis of the literature, the role of its corresponding protein in resistance, virulence, metal transport, or biofilm formation is indicated.

**Figure S4.**
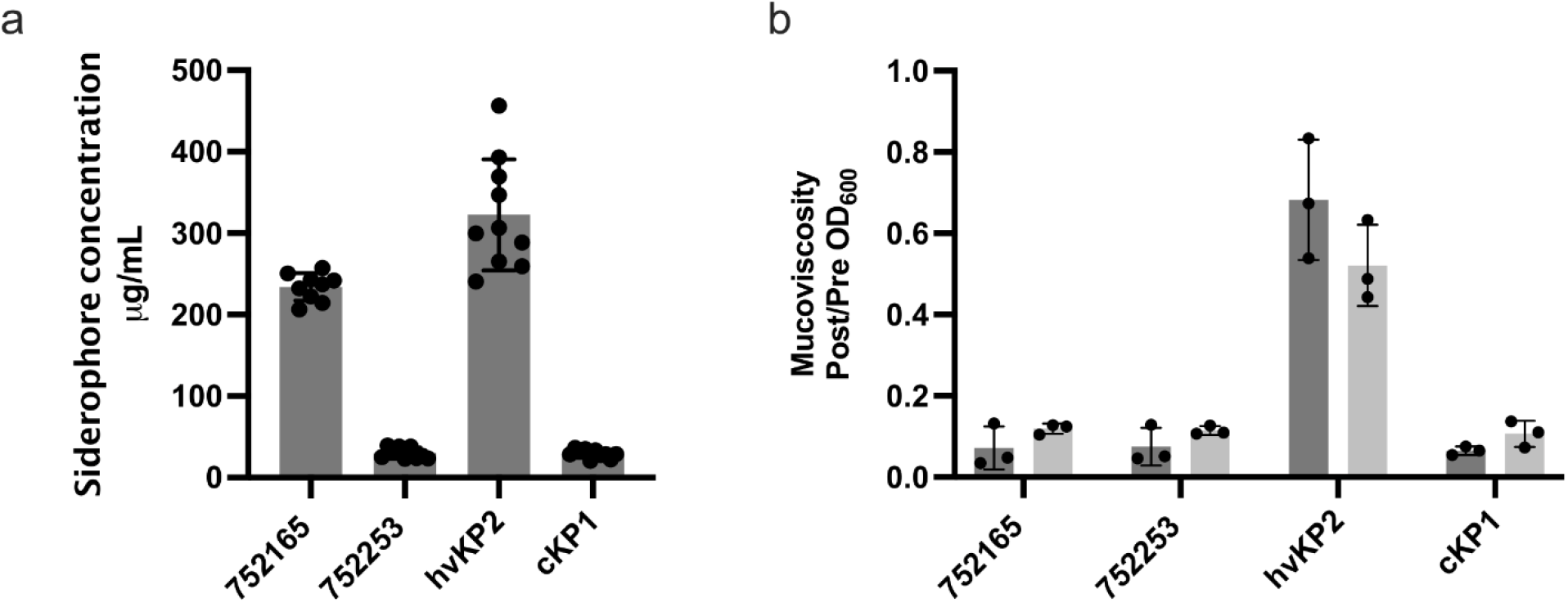
**(A)** Quantitative siderophore production of outbreak isolates 752165 (pSI0739-ARMA-Vir) and 752253 (pSI0739-ARMA-Vir, event B). **(B)** Mean mucoviscosity of outbreak isolates 752165 and 752253 compared to reference *K. pneumoniae* isolates. Independent assays were performed for both media (LB and M9). Reference hypervirulent (hvKP2) and “classical” (cKP1) isolates are shown throughout all panels for comparison.

**Fig. S5.**
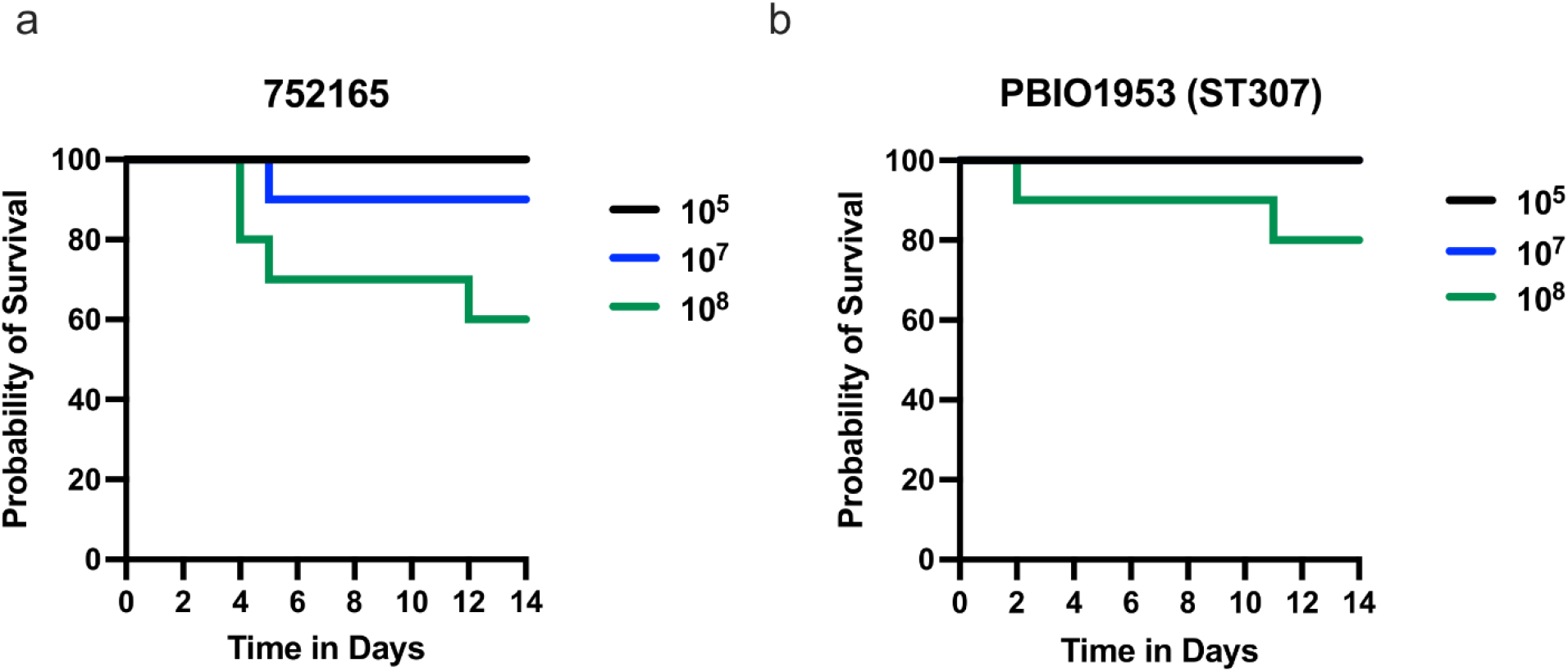
Kaplan-Meier survival curves of outbred CD1 mice after subcutaneous (SQ) challenge with 10^5^, 10^7^, or 10^8^ CFU of **(A)** outbreak isolate 752165 and (**B**) ST307 strain PBIO1953. Total *n*=10 (*n*=5 in each of 2 independent experiments) for each titer for each strain.

